# Stimulating neurogenesis of distinct retinal lineages in human retinal pigmented epithelium (RPE) with proneural transcription factors

**DOI:** 10.64898/2026.06.17.732991

**Authors:** L. Kaplan, A.L. Green, J. Pang, M. Pavlou, J. Wohlschlegel, T.A. Reh

## Abstract

There are currently few promising approaches for treatment of photoreceptor pathologies: for example, gene therapy to augment or replace mutated genes, has proven successful in preclinical studies, and some of these therapies are moving towards the clinic. Another approach aims to unlock the inherent stem-cell potential of non-neuronal retinal cells to regenerate neurons in situ. This line of research is based on the discovery that some vertebrates can restore even severely damaged retina from RPE with all the necessary cell types to regain full functionality.

To determine whether this approach can be applied to humans, we established a robust in vitro culture system using fetal human RPE, and employed a barcode-multiplexed, single cell RNAseq based screen to find factors that would reprogram human RPE into photoreceptors.

With this approach we were able to identify NEUROD1 as a complimentary factor to ASCL1. Their combined overexpression together with a treatment with bFGF and Activin A inhibitor produced RPE-derived neuronal cells with expression patterns characteristic of photoreceptors and other lineages.

## Introduction

Retinitis pigmentosa (RP) affects approximately 1.5 million people worldwide [1]. It causes major and irreversible loss of, initially, rod photoreceptors in the periphery and eventually leads to cone death and blindness. Current developments in gene therapy, which aims to deliver healthy copies of mutated genes, show promising results and are increasingly becoming available in the clinic. [2], [3]. Other methods, such as retinal prosthetics and optogenetics, rely on the fact that the inner retina, harboring interneurons and Müller glia, remains relatively intact even at late disease stages [4], [5], [6].

Ideally, the best therapy would be to replace the photoreceptor cells lost to RP with new rods and cones. This concept is being explored in the form of the differentiation of pluripotent stem cells into functional neurons and their implantation into affected retina [7]. A similar approach aims to unlock the inherent stem-cell potential of non-neuronal retinal cells to regenerate neurons *in situ.* This line of research is based on the discovery that some vertebrates can restore even severely damaged retina and retinal pigment epithelium (RPE) with all the necessary cell types to regain full functionality [8], [9], [10].

The source of regenerated retinal cells varies between species: in amphibians the regenerated retina arises primarily from the RPE, while in fish the new neurons are regenerated from Müller glia [8], [9], [10]. Using the knowledge of genes involved in this process, there have been successful efforts by our lab and others to reactivate these capabilities in mammalian Müller cells and reprogram them to neurons of the inner retina, like bipolar or ganglion cells, using proneural transcription factors, such as Ascl1, Atoh1, Pou4f2 and Islet1 [11], [12]. While several reports have documented regeneration of functional inner retinal neurons in the last ten years, it has been more difficult to stimulate robust regeneration of photoreceptors from Müller glia in mammals.

Although it has been known for some time that RPE in amphibians has the capacity for retinal regeneration, relatively few studies have attempted to stimulate this process in mammals. Prior attempts to stimulate neural regeneration from RPE in species that do not normally regenerate retina, have met with some success. In developing chick embryos for example, FGF stimulates retinal regeneration at embryonic day 3.5, and antagonizing the activin receptor can extend the window of opportunity when FGF is effective [13]. Infection of developing chick or immortalized human RPE with proneural transcription factors, such as Neurod1 or Neurogenin, can shift RPE towards a neuronal fate that resembles photoreceptors [14], [15].

To advance this approach in a more physiologically relevant model, in this study we show that acutely isolated fetal human RPE can be reprogramed into various neuronal subtypes. Overexpressing proneural factor ASCL1 activates a neuronal expression pattern similar to reprogrammed human Müller glia. Given the clinical need for regenerating photoreceptors, we employed a barcode-multiplexed, single cell RNAseq based screen for factors that would push human RPE towards a photoreceptor fate. With this approach we were able to identify NEUROD1 as a complimentary factor that not only increased the reprogramming efficiency but also yielded cells with photoreceptor characteristics from primary human RPE.

## Results

### Establishing primary human RPE cultures

We first aimed to establish and characterize an *in vitro* system to reliably culture primary human RPE and verify it as an appropriate testbed for subsequent reprogramming experiments. Human fetal RPE was peeled in coherent sheets from dissected eyes (ages ranging around 70-150 days of gestation) and enzymatically digested to derive small cell clumps. These were seeded on Matrigel coated coverslips and cultured for up to 30 days. The colonies attached and proceeded to proliferate until a homogeneous monolayer of hexagonal cells was achieved (Fig. 1A). We monitored proliferation using MKI67 staining, which revealed a peak around 8 days *in vitro* (DIV) and a subsequent decrease until approximately 20 DIV at which point almost no MKI67 positive cells were detected and the cell density reached a plateau (Fig. 1B). Immunofluorescence staining for canonical RPE markers ZO1, PMEL and OTX2, as well as pigmentation onset, confirmed a protein expression profile that was representative of *bona fide* RPE (Fig. 1C). The cell-membrane associated ZO1 staining allowed us to use the machine learning based software REShAPE [16] to compare several morphological features between native flat-mounted and cultured human RPE across thousands of individual cells in an unbiased manner. We found no difference in the number of neighbors and hexagonality between native and cultured fetal human RPE, though there was a moderately increased size in the latter (Fig. 1D).

**Figure 1:**
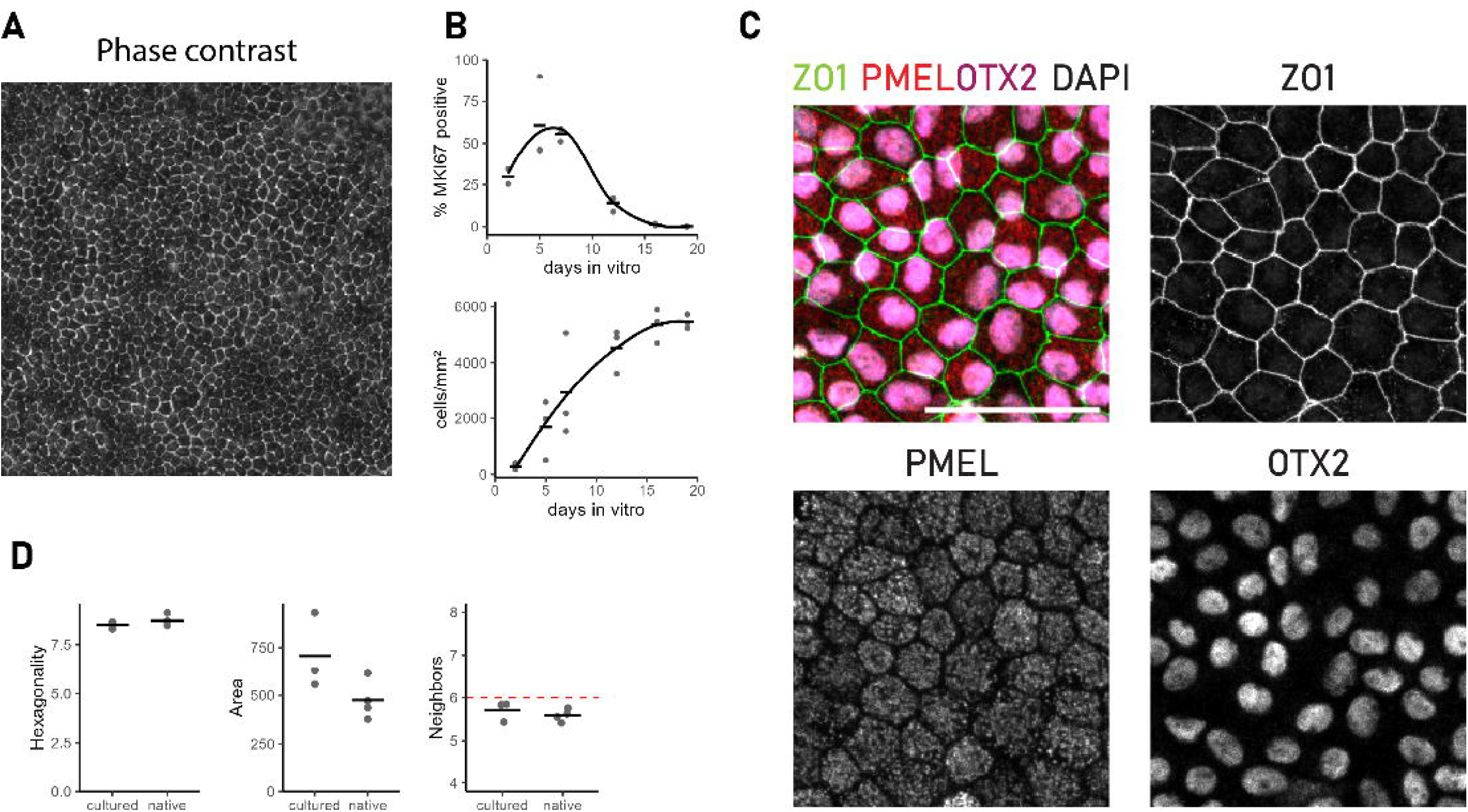
A) Phase contrast image of human fetal RPE harvested from a donor eye of gestation day 137 (D137) and cultured for 22 days in vitro (DIV). B) MKI67 staining of cultured RPE that was fixed at different timepoints revealed peak proliferation around 8 days in vitro (DIV8). Proliferation ceased and cell density reached its maximum at around DIV13. n =3, ages: D113, D115, D126. Horizontal bars indicate mean. C) D130 RPE was cultured until DIV20 and stained for RPE markers ZO1, OTX2 and PMEL Scale bar 50 µm. D) ZO1 staining from confluent RPE or native fetal RPE wholemounts was used for segmentation of outlines of individual cells via REShAPE [16]. Cultured RPE showed a slight increase in cell area but no difference in the number of neighbors or hexagonality. cultured: n=3, ages: D78, D130, D137. native: n = 4, ages D72, D78, D85, D145. Black bars indicate the mean and dashed red line indicates 6 neighbors.

### Strong expression of ASCL1 induces neuronal expression pattern in human RPE

As a proof of principle we wanted to see whether ASCL1 overexpression alone is capable of driving RPE-to-neuron reprogramming in a similar manner as previously described for fetal human Müller glia [11], [17]. We infected cultured fetal RPE with lentiviral vectors each carrying an ASCL1-GFP overexpression cassette either under a strong ubiquitous promoter (*CMV>ASCL1-TuGFP*) or the glial-specific HES1 promoter (*pHES1>ASCL1-eGFP*) [17] (Fig. 2A), as previously published scRNAseq data of human RPE revealed HES1 expression [18]. After 8 days, we recorded ASCL1 protein expression in GFP positive cells with both vectors, confirming successful transduction and the activity of the HES1 promoter in RPE (Fig. 2B). Additionally, some cells infected with *CMV>ASCL1-TuGFP* exhibited axon-like protrusions that were positive for TUBB3 (TUJ), indicating a shift to a neuronal fate. We did not encounter this morphology where ASCL1 was driven off of the HES1 promoter (Fig. 2B).

**Figure 2:**
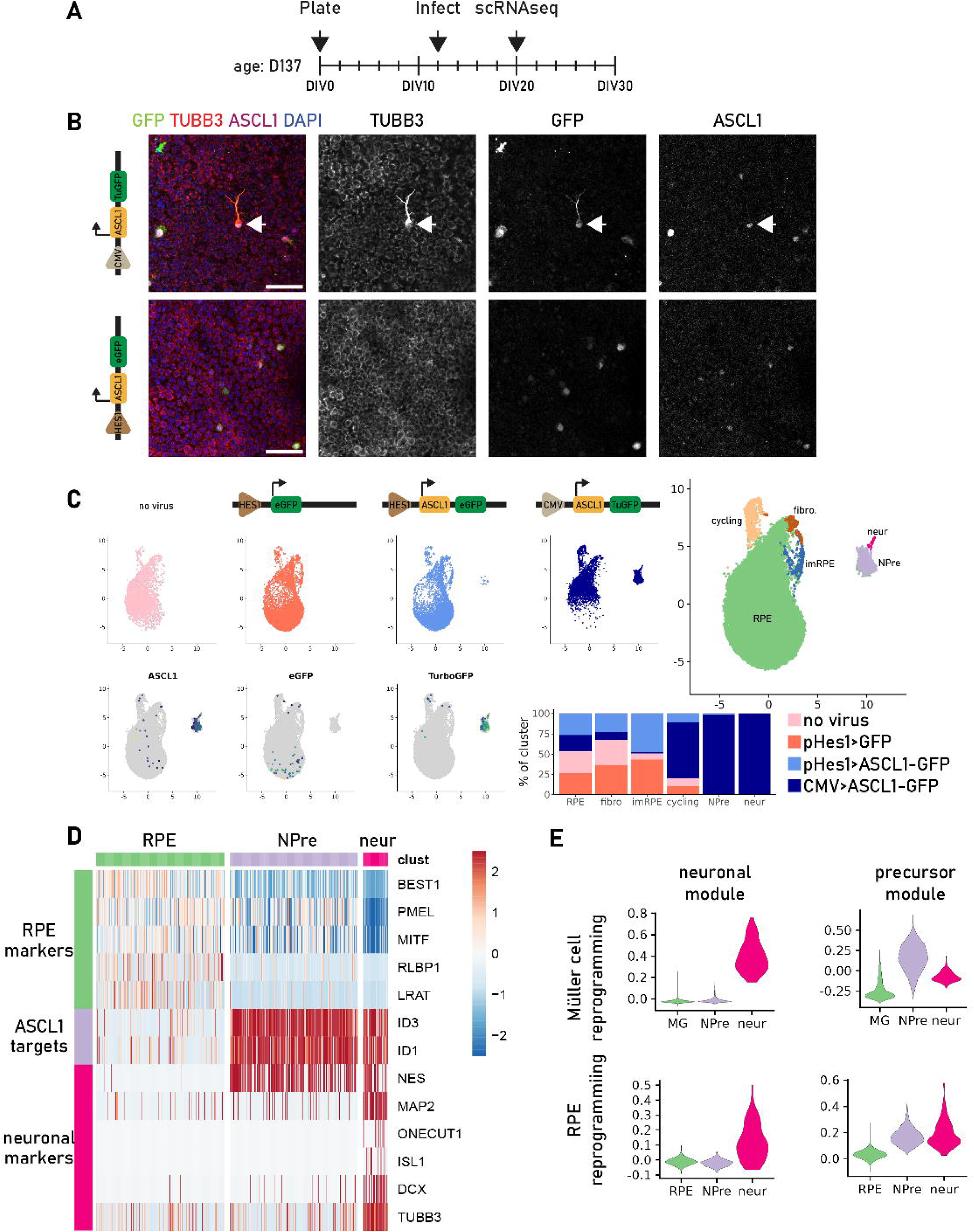
A) Overview of scRNAseq experimental timeline. Cultured human RPE of a D137 donor was infected at DIV12 with a lentivirus carrying either a CMV>ASCL1-TuGFP, pHES1>ASCL1-eGFP or pHES1>eGFP cassette; one sample was left uninfected. Cells were harvested at DIV20 and 8 days post infection (DPI). B) CMV and Hes1 promoter were both capable of driving the expression of transgenes producing GFP/ASCL1 double positive cells. Only CMV>ASCL1-TuGFP produced cells with increased TUBB3 (TUJ) expression and axon-like morphology (white arrow). Scale bar 50 µm C) ScRNAseq analysis revealed a big cluster of cells with equal contributions from all samples (RPE). Two clusters contained exclusively cells that were transduced with ASCL1 carrying virus and were dominated by CMV>ASCL1-TuGFP expressing cells (NPre, neur). Three clusters showed expression patterns characteristic for proliferating cells (cycling), fibroblasts (fibro) and subconfluent RPE (imRPE). D) NPre cells showed a downregulation in RPE markers and an upregulation of general neuronal genes, as well as ASCL1 targets. Neur exhibited mature neuronal expression pattern. E) From a previously published dataset in which fetal human Müller glia were reprogrammed with ASCL1, we identified genes characteristic for NPre and neur. We compared their combined expression in the form of module scores between Müller glia and RPE derived reprogrammed cells. Genes that were induced upon ASCL1 overexpression in Müller cells and yielded NPre (precursor module) or neur (neuronal module), were also upregulated in the respective clusters in ASCL1-transduced RPE.

To elucidate the extent of reprogramming and whether distinct cell types were produced, we performed single cell RNA sequencing (scRNAseq) on RPE cultures transduced with lentiviral vectors carrying either *CMV>ASCL1-TuGFP*, *pHES1>ASCL1-eGFP* or *pHES1>eGFP* cassettes; we used an untransduced control sample as a baseline (Fig. 2C). The majority of cells from all samples formed a single cluster expressing RPE genes, such as BEST1, RLBP1 and LRAT (Fig. 2C, 2D). Two clusters that contained equal numbers of cells originating from samples with and without ASCL1, expressed fibroblast-related genes (Fig. 2C). These likely originate from a population of fibroblasts that was carried over during dissection. The cluster we termed immature RPE (imRPE) expressed both RPE markers and genes associated with epithelial to mesenchymal transition (EMT, GO:0001837), indicating they might be subconfluent RPE. (Suppl. Fig. 1A). The cluster of cycling cells was comprised mostly of cells from the *CMV>ASCL1-TuGFP* sample, though all conditions had dividing cells.

Overexpression of ASCL1 produced two clusters with expression of genes normally found in neuronal precursors (Npre) or more mature neurons (neur); cells in these clusters originated almost exclusively from *CMV>ASCL1-TuGFP*. Residual expression of RPE markers and a lack of fibroblastic genes indicated that they likely arose from RPE rather than fibroblasts (Suppl. Fig. 1A). Differential gene expression analysis of these two clusters revealed both (1) the downregulation of RPE genes such as TTR and LRAT and (2) the enrichment of neuronal genes such as NES and DSCAM, as well as ASCL1 downstream targets ID1/3 and DLL3 (Fig. 2B, Suppl. Fig. 1B). The larger had higher expression of these neural progenitors, while the smaller cluster expressed more lineage-specific markers, such as ONECUT1, ISL1 and TH; this pattern of gene expression suggests that ASCL1 directs RPE reprogramming towards an amacrine/horizontal cell-like fate (Fig. 2D).

Next, we compared the expression changes that resulted from ASCL1 overexpression between Müller cells and RPE. For this we reanalyzed scRNAseq data from a previous publication where human fetal Müller cells were cultured in 2D and infected with *CMV>ASCL1-GFP* [11]. For a more global and unbiased approach we defined the top 100 differentially expressed genes between Müller cells and NPre or neurons (Suppl. Table 1) and combined them into two module scores that captured the transcriptomic profile of the respective clusters. As expected, the neuronal score was highest in neurons while the precursor score was elevated in cells of the NPre cluster and to a lower degree in neurons. Subsequently, we used the same gene sets to generate scores for the analogous clusters from the reprogrammed RPE (Fig. 2E). Here, the newly generated neurons elicited a similar expression pattern: they showed a specifically increased neuronal score as compared to RPE or NPre. Interestingly, although the precursor score was lowest in RPE, both neurons and NPre showed similar values. This analysis indicates that ASCL1 overexpression drives neuronal reprogramming and activates gene regulatory networks that produce comparable results in both human fetal Müller glia and

### Multiplexed screen for the identification of pro-photoreceptor genes

While the overexpression of ASCL1 in RPE seemed to produce neuron-like cells with some markers of mature inner retinal neurons, our goal is to produce photoreceptors. As noted above, previous reports used proneural transcription factors to stimulate production of photoreceptor-like cells from developing chick RPE or immortalized RPE cell lines [14], [15], [19], [20]. To find more potential candidate genes, we mined several publicly available and in-house datasets representing human retinal development (see the methods section, Suppl. Fig. 2). In brief, we used single cell chromatin accessibility and transcriptomics [11], [21], [22], [23] to find pseudotime trajectories leading from progenitor cells to photoreceptors. We then identified genes whose expression, open chromatin or binding motif accessibility follow this trajectory and importantly, do not lead to the bipolar cell fate (Suppl. Fig. 2A). Some notable examples include transcription factors like RAX2, LMO1 and NEUROD1, but also genes with other functions like AIPL1, which is thought to have chaperone activity [24]. Another source of candidate genes was a transcriptome dataset from FGF-mediated reprogramming of embryonic chicken RPE [25], from which we identified genes that are specifically activated during that process (Suppl. Fig. 2B). Additionally, we were interested in the role of OTX2 during retinal development as it is expressed in photoreceptors, bipolar cells and also in RPE. To explore which regulatory networks OTX2 is driving in these different cell types, we compared bulk chromatin accessibility from fetal RPE [26] with pseudobulk data derived from bipolar cell and photoreceptor clusters from single cell ATACseq [23]. Specifically, we looked for OTX2 binding motif-containing regions that were accessible in photoreceptors and RPE but not in bipolar cells (Suppl Fig. 2C). Using transcription factor binding site analysis on these peaks revealed an enrichment of members of the ETS family like ELK1/4 and ETV1/4.

From our list of candidates, we picked 5 promising genes that were also reported to play a potential role during retinal development (LMO1, NEUROD1, RAX2, EZH2, ETV1). As we aimed to amplify the neurogenic potential of ASCL1, we created combinations with our candidates. Furthermore, we wanted to test the neurogenic potential of some novel genes and compare it to ASCL1 alone which resulted in a lentiviral library of 7 constructs: ASCL1, ASCL1-EZH2, ASCL1-LMO1, ASCL1-NEUROD1, ASCL1-RAX2, NEUROD1-ETV1, RAX2.

Our previous experiment showed that RPE-derived neuron-like cells can be identified and characterized as newly generated clusters in single cell transcriptomics. In order to test the effects of all reprogramming candidates at once using scRNAseq, we designed a lentiviral vector system that allows the simultaneous expression of a transgene cassette, a fluorescent reporter, and a construct-specific barcode. The transgene and barcode are expressed from the same locus on the antisense strand with its own transcription termination signal (Fig. 3A). This is necessary to allow the barcode to be in direct proximity to a polyadenylation site without the internal disruption of the transcription of the viral genome. Taken together this increases the probability for a barcode to be captured and detected in a standard 3’ biased droplet based scRNAseq protocol. Furthermore, having a fluorescent reporter that is driven off a distinct ubiquitous promoter facilitates (1) an internal control of vector expression and (2) positive selection of transduced cells independently of a reprogramming outcome.

**Figure 3:**
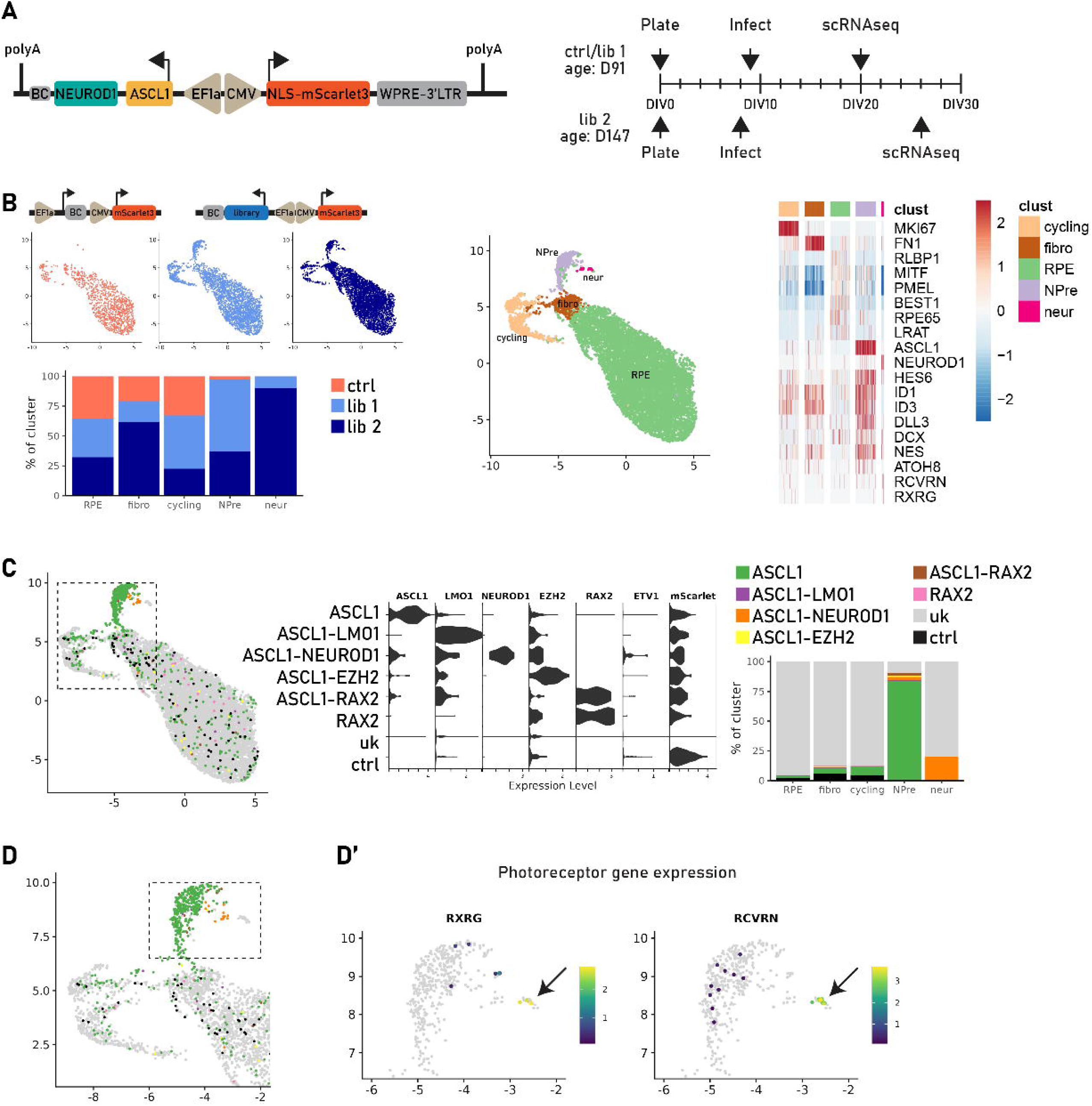
A) Example structure of constructs present in the screen library. CMV drives the expression of fluorescent reporter mScarlet3. Adjacent but in antisense direction, EF1a promotor regulates the expression of the transgene cassette. It contains the sequences of the candidate genes followed by a construct-specific barcode (BC) and a polyadenylation signal (polyA). B) Human fetal RPE was derived from two donors aged D91 (ctrl/lib 1) and D147 (lib 2) and infected during peak proliferation (DIV8/DIV9). Cells were harvested at DPI11 (ctrl, lib 1) or DPI18 (lib 2) for scRNAseq. B) UMAP embedding revealed one big cluster with equal contributions from all samples and was determined to be RPE. Cycling cells and fibroblastic (fibro) cells were also present in all samples. Only cells transduced with the library formed new clusters NPre and neur. These showed increased expression of ASCL1, its downstream targets, and several neuronal markers concomitant with a downregulation of RPE markers. C) Cells were assigned to the construct they were transduced with by the detection of its specific barcode. Transgene expression matched the assigned construct. Cells with unknown barcode (uk) showed no specific upregulation of transgenes or the fluorescent protein while those that were assigned to the control vector (ctrl), exhibited strong expression of mScarlet. Most of the cells of NPre expressed the ASCL1 construct, while neur seemed to arise from the combination of ASCL1 and NEUROD1 overexpression. D) Zoomed in UMAP focusing on the cluster containing reprogrammed cells. Dashed box shows cells presented in D’. D’) black arrow: Cells in the neur cluster expressed photoreceptor genes RXRG and RCVRN.

We infected RPE cultures from two donors with the same library and performed scRNAseq at 11 or 18 days post infection (DPI), to allow for a longer maturation time (Fig. 3A). Again, we saw a large cluster with equal contributions from all samples expressing RPE markers, a smaller cluster of cycling cells and a cluster with some EMT/fibroblast gene expression (Fig. 3B, Suppl. Fig. 2D, 2E). However, only cells originating from samples infected with the lentiviral library produced two new clusters (NPre and neur) and their expression pattern confirmed their RPE origin (Suppl. Fig. 2B).

We then used the barcodes to assign each cell to the construct it was transduced with and confirmed the expression of the respective genes (Fig. 3C). While the cell-construct mapping was relatively sparse, and we did not find any cells infected with NEUROD1-ETV1, we were able to assign a majority of the cells in the newly generated cluster NPre. This suggests that if transgene expression is strong enough for reprogramming it is also more likely to produce enough barcode for demultiplexing. The NPre cluster consisted mostly of cells expressing either ASCL1 alone or in combination with other genes, confirming its central role in the reprogramming process (Fig. 3C). The only condition that produced a new neuronal cluster (neur), that was different from ASCL1 alone, was the combination ASCL1-NEUROD1 (Fig. 3D). Cells in this cluster showed expression of RXRG and RCVRN indicative of a photoreceptor identity. No cells with this gene profile were identified in any other scRNAseq sample, indicating that carryover of native photoreceptors is unlikely.

### ASCL1-NEUROD1 reprograms RPE into neurons of different retinal lineages

Our screen showed that the addition of NEUROD1 had the potential to shift the fate of reprogrammed RPE to a photoreceptor-like state. However, this process was very inefficient as it produced only a small number of neuronal cells compared to even ASCL1 alone. To improve on this potential, we generated a new expression vector that (1) simplifies the cassette and (2) replaces the EF1a with the CMV promoter (Fig. 4A), as the latter proved to drive stronger expression in human fetal RPE cultures in our hands. Additionally, we employed a treatment with bFGF and activin A inhibitor (SB431542) which was shown to promote RPE to retinal reprogramming in the chicken retina [13], [25], [27] (Fig. 4A). For this paradigm, we transduced cells at their proliferation peak (8 DIV) (Fig. 1B) and started bFGF + SB431542 treatment (FA) two days later to allow for vector expression onset. We included EdU in the culture media at 13 DIV, when RPE proliferation plateaus (Fig. 1B), to track newly generated neurons without oversampling for cycling RPE (Fig. 4A). Although cells with neuronal marker expression were present at earlier timepoints (Suppl. Fig. 3A), we chose to harvest cells at 20 DPI to allow for longer maturation. We collected mScarlet+ transduced cells using fluorescence-activated cell sorting (FACS) and performed scRNAseq or immunostaining. As in the previous experiments, a cluster made up of equal parts from all samples appeared and represented unaltered RPE. The overexpression of ASCL1-NEUROD1 produced 5 new clusters (reprog 1-5, Fig. 4A, 4B). Reprog 1 was comprised mainly of cells treated with ASCL1-NEUROD1 but no FA supplement, whereas reprog 2-5 were cells treated with ASCL1-NEUROD1 + FA (Fig. 4B). As expected at 28 DIV, there were only few cells remaining in a proliferative state. However, their proportion was markedly increased in the FA treated sample, likely due to the effect of bFGF on proliferation [28], [29]. Expression of ASCL1-NEUROD1 led to the downregulation of RPE genes like LRAT, RPE65 and BEST1 and an increase of ASCL1 downstream targets HES1, ID1/3 and DLL3 (Fig. 4C). Indeed, we found expression of marker genes of cell types emerging at different stages during retinogenesis: amacrine cell markers TFAP2A and TFAP2B [30], [31] were specifically expressed in reprog 3 and ganglion cell marker EBF1 [32] was found across reprog 3 through 5 (Fig. 4C). Most strikingly, reprog 4 cells expressed markers of progenitor cells (VSX2), bipolar cell subtypes (FEZF2, VSX1, PRDM8), but also photoreceptors (RCVRN, NRL, AIPL1) and ganglion cells (ELAVL3) (Fig. 4C).

**Figure 4:**
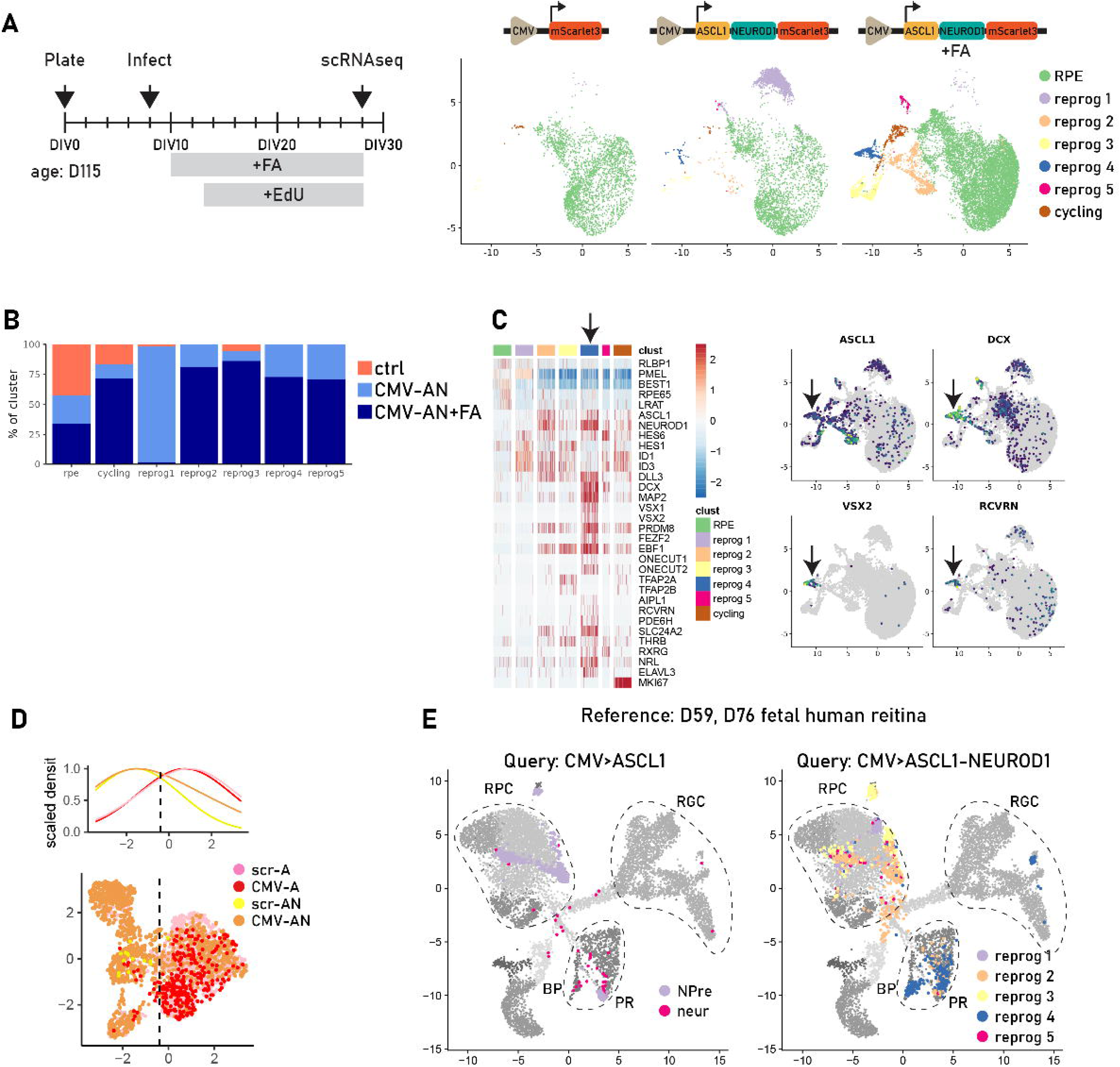
A) RPE from a D115 human donor was cultured for 8 days before infection. At DIV10 bFGF and Activin A inhibitor (FA) were added to the media and on DIV13, when proliferation declined, EdU application was started. Cells treated with control virus, with CMV>ASCL1-NEUROD1 or CMV>ASCL1-NEUROD1 plus FA were harvested on DPI 20 and processed for FACS enrichment and scRNAseq. Only samples that received ASCL1-NEUROD1 overexpression generated new clusters and FA treatment increased their proportion. B) The RPE cluster was equally represented by all samples, while there was an enrichment in FA treated cells in the cycling cluster. The other clusters were dominated by cells originating from samples infected with the transgene virus. C) Residual but markedly downregulated markers of RPE were found in all reprog clusters. ASCL1 and NEUROD1 overexpression activated downstream targets like ID1/3 and DLL3 as well as general neuronal markers like DCX and MAP2. Reprogrammed cluster 4 (Reprog 4, black arrow) was enriched in mature neuronal markers of different lineages like VSX1/VSX2 (bipolar cells), ELAVL3 (ganglion cells), NRL (rods), RCVRN (photoreceptors). D) ScRNAseq data from CMV>ASCL1 (CMV-A), EF1a>ACSL1 (scr-A), CMV>ASCL-NEUROD1 (CMV-AN) and EF1a>ASCL1-NEUROD1 (scr-AN) from previous experiments were integrated and coembedded in a UMAP. While there is significant overlap between single and dual transcription factor overexpression, the addition of NEUROD1 clearly shifts the resulting cell fate. Top plot represents the scaled density of cells along the first UMAP axis. E) Projection of reprogrammed RPE treated either with CMV>ASCL1 (left) or CMV>ASCL1-NEUROD1 (right) onto a combined reference UMAP from D59 and D76 human fetal retina. ASCL1 overexpression alone produces cells that align with retinal progenitors (RPC), bipolar cells (BP) and photoreceptors (PR). The addition of NEUROD1 produces a higher proportion of bipolar cells and photoreceptors (mostly from reprog 2 and 4) but also shows prominent subclusters mapping to native retinal ganglion cells (RGC) and the transition point between RPC to mature neurons.

To gauge the influence of the addition of NEUROD1, we integrated cells expressing ASCL1 or ASCL1-NEUROD1 from our previous run (Fig. 3A) with the reprog clusters of our optimized paradigm (Fig. 4A). The resulting UMAP embedding revealed a general overlap of expression patterns between cells treated with ASCL1 alone or ASCL1-NEUROD1. Across all experiments, the addition of NEUROD1 yielded reprogrammed neurons with a more mature transcriptomic profile (Fig. 4D). To characterize these neurons in an unbiased manner, we used a human fetal scRNAseq data set [11] as a reference (Suppl. Fig. 3B) and projected the ASCL1 or ASCL1-NEUROD1 reprogrammed cells onto its UMAP space. Cells reprogrammed with ASCL1 alone overlapped mostly with retinal progenitors and some with photoreceptors and bipolar cells. The addition of NEUROD1 yielded a similar mapping in the progenitor cell clusters but now also a substantially increased coverage in the bipolar cells and cone clusters, as well as a few cells in the developing ganglion cell branch (Fig. 4E).

### Clusters of reprogrammed RPE form domes containing cells of different lineages

The single cell analysis suggested that our reprogramming paradigm generates cells with a transcriptomic profile that resembles developing retinal neurons of distinct lineages. To characterize these cells further, we performed EdU detection and immunostaining for the general neuronal marker DCX, found in reprog 1/2/4/5 (Fig. 5A). FA-treated cells that were transduced with *CMV>ASCL1-NEUROD1-mScarlet*3 showed prominent mScarlet+ clusters that comprised cells protruding from the plane of the surrounding RPE monolayer in a rosette like formation (Fig. 5A,B; Suppl. Fig. 4A). In addition to their 3D organization, most of these clusters were positive for DCX revealing a complex network of axonal outgrowths (Fig. 5A). However, there were only few cells with strong EdU signal (Fig. 5A), indicating that most of them were born before the start of the EdU addition (i.e. <DIV13, <DPI5) and did not proliferate further. Without FA treatment we found similar DCX/mScarlet positive clusters, albeit fewer and with shorter arbors (Suppl. Fig. 4B, top). Infection with the control virus carrying only mScarlet did not produce clusters or DCX positive cells (Suppl. Fig. 4B, bottom). The single cell transcriptomic analysis of the reprog 4 cluster suggested a generation of HuC/D (ELAVL3/4) and VSX1 expressing cells in distinct subpopulations. Immunostaining showed cells inside the rosette-like structures were positive for HuC/D and arranged in a dome shape, suggesting the formation of polarization or lamination (Fig. 5C). VSX1 and HuC/D staining were mutually exclusive but often occurred side by side in the same cell cluster (Fig. 5C). VSX2 and RCVRN staining revealed a heterogeneous composition of the neurogenic clusters. Some contained VSX2 positive cells without any RCVRN staining (Fig. 5D top row), while others seemed to have cells expressing either of the markers (Fig. 5D center row). Strikingly, some larger colonies of reprogrammed cells had extensive arborization of RCVRN positive processes (Fig. 5D bottom row). Overall, we were able to validate human RPE reprogramming into distinct neuronal classes using both lineage-specific transcripts detected via scRNAseq and immunocytochemistry of their respective proteins.

**Figure 5:**
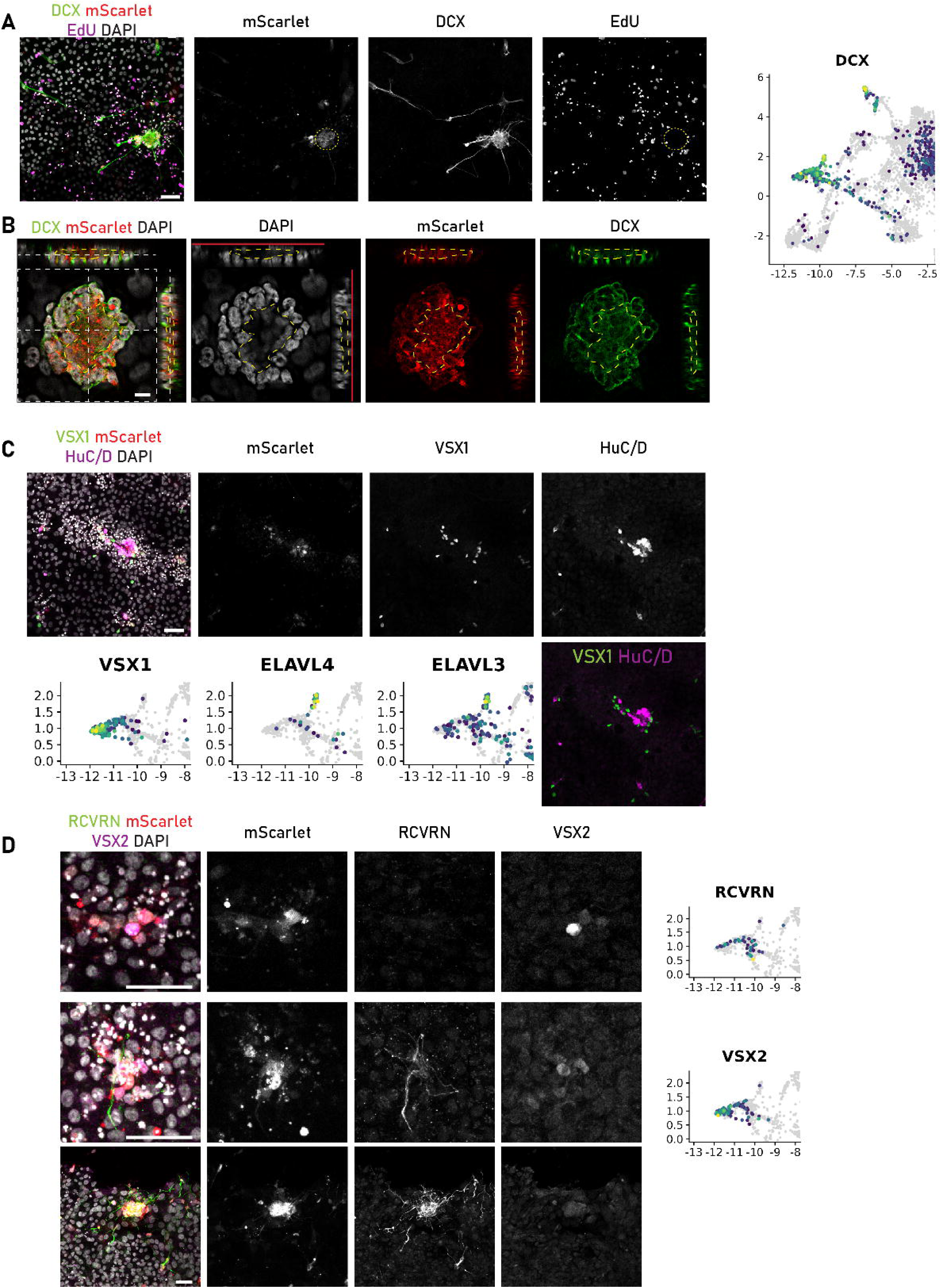
Cultured RPE from the same donor and experimental run that was used for scRNAseq (Fig. 4) were processed for immunostaining. All images that are shown here are from the CMV>ASCL1-NEUROD1+FA paradigm. A) Transduced cells, represented by mScarlet signal, were accumulated in clusters and exhibited distinct DCX positive axon-like arborization and were surrounded by EdU positive, proliferating RPE. Reprogrammed cells were not positive for EdU (dashed yellow ellipse). B) Orthogonal section view of a neurogenic dome produced by CMV>ASCL1-NEUROD1 and FA treatment (DPI20) of RPE derived from a D110 donor. Dashed, yellow outlines depict the location of the hollow inner space. Red lines indicate the location of the glass coverslip that the cells were grown on. The dome appears flattened due to being compressed between the coverslip and the microscope slide. Right: DCX transcript expression zoomed on reprogrammed clusters. C) Some clusters contained cells that were positive for either HuC/D or VSX1 in a mutually exclusive manner confirming the transcriptomic pattern of reprog 4 (Fig. 4). D) VSX2 and RCVRN expression was heterogeneous, sometimes present in the same cluster (center row) or one being expressed without the other (top row: only VSX2; bottom row: only RCVRN). Scale bar A, C, D: 50 µm. Scale bar B: 10 µm

## Discussion

Loss of retinal neurons, is currently an irreversible process and although therapies exist to treat some pathologies [2], [33], [34], they mostly just halt or decelerate further degeneration. No therapy exists at present that is capable of regenerating lost neurons in the adult retina or other areas of the central nervous system. For successful restoration of vision, it is therefore crucial to develop methods for replacement of retinal neurons, particularly photoreceptors. In nonmammalian vertebrates, natural photoreceptor regeneration occurs after injury using two endogenous cells sources: Müller glia (MG) and the RPE. After injury to the retina, MG undergo a spontaneous reprogramming, expressing progenitor genes and generating new neurons in birds, fish and frogs. Fewer animals utilize the RPE to regenerate retina; this source is primarily used in urodele amphibians, larval anurans, and embryonic chick. We have previously shown that retinal regeneration can be stimulated in mammals, using both transgenic models or a combination of viral vectors and transgenic lineage tracers [35], [36]. The overexpression of ASCL1 alone, along with the histone deacetylase inhibitor Trichostatin-A, can trigger MG reprogramming into progenitor-like cells and neurons after injury in mice [37]. Further combination with Atoh1, Pou4f2 or Isl1 tunes the reprogramming efficiency and increases the diversity of neuron classes generated by the MG [35], [38]. Similar approaches have also identified the Notch and nuclear factor I pathways as inhibitory for mammalian neural regeneration from MG [39]. Therefore, it would seem that a relatively small number of factors limit regeneration from MG in mammals and further developments in this area may well lead to a regenerative therapy for humans.

By contrast with the progress in stimulating neurogenesis from MG in mammals, there have been fewer attempts to stimulate regeneration from the RPE in mammals. This is surprising, since their regenerative capacity in amphibians and embryonic chick is remarkable and can regenerate entire laminated retina. The regenerative capacity of human RPE remains unclear, which motivated our efforts to further develop the RPE as an accessible platform for testing reprogramming paradigms. We were able to extract primary human fetal RPE that readily proliferated until confluency and achieved characteristic morphology and gene expression relevant to native human tissue. We then tested the neurogenic potential of human RPE cells after ASCL1 overexpression and found that RPE can indeed be coaxed to produce neuron-like cells with an expression pattern similar to those of Müller glial origin.

Attempts to stimulate regeneration from the RPE in birds and mammals have focused on two approaches: treatment with signaling molecules such as FGF and viral overexpression of proneural factors, such as Neurogenin. For example, treatment with bFGF was shown to induce RPE-to-retina transdifferentiation of embryonic chick and rat retina, a process that was promoted by inhibiting Activin A signaling and DNA demethylation [13], [20], [27], [40], [41]. Promising results have also been obtained in chick embryo RPE with viral expression of neurogenins and neuroD, which expressed photoreceptor markers and in some cases seemed to be responding to light [14], [19]. NeuroD1 specifically, seemed to also be effective in initiating the transcription of photoreceptor genes in mammalian cells [15]. More recently, a multistep paradigm involving overexpression of ASCL1 and miR-9/9*-124 combined with the knockdown of PTBP2 and p53 was established for reprogramming immortalized RPE-like cells [42]. However, this study had some limitations: While the treatment clearly seems to produce neuron-like cells positive for general markers MAP2 and TUJ, the authors also claim a high percentage of reprogrammed cells express photoreceptor markers. Although immunofluorescence staining suggested nearly all reprogrammed cells are positive for both rod and cone opsin, this was not reflected in their transcriptomic analysis.

Drawing from the lessons learned from these studies we first sought to find a suitable transcription factor combination and then enhance it with the FGF plus Activin A inhibition treatment. Our screen for proneural genes confirmed that ASCL1 alone is capable of driving RPE reprogramming into cells with neuronal characteristics. In our scRNAseq analysis of RPE reprogramming, we found that the cells infected with ASCL1 alone showed the greatest degree of neurogenesis. It is not clear why the other factors in the screen were less effective. From previous studies, we know that ASCL1 has the potential to induce proliferation, and so this may have resulted in an expansion of the ASCL1-expressing RPE [37]. On the other hand, viability of mature neurons in culture without specific media is limited, so if any other vector produced cells that died before harvesting, they would not appear in the data. Finally, a preferential virus generation from the plasmid library cannot be ruled out and insufficient ASCL1 expression in the combinations might have led to a lack of reprogramming, leading to false-negatives. Importantly, there was no newly generated cluster without any assigned reprogramming construct, and only ASCL1-NEUROD1 produced a small cluster with photoreceptor-like expression outside of the Npre cluster. However, it is important to note that the subcluster with the highest level of RCVRN and RXRG was not assigned to an ASCL1-NEUROD1 identity by their barcode, though they express detectable levels of NEUROD1 and are only present in the lib2 sample (Figure 3B).

The overexpression of ASCL1 and NEUROD1 produced cells with neuronal morphology and expression of general neuronal markers and the treatment with FA vastly enhanced this process, as evident from an increase in both proliferating and reprogrammed cell clusters concomitant with expression of mature neuronal markers. From the single cell data we saw that one cluster seemed to consist of cells expressing mature neuronal genes of different developmental branches. Initially, we assumed that individual reprogrammed RPE cells might result in different neurons. Surprisingly, immunostaining revealed that reprogrammed cells seemed to be arranged in neurogenic domes protruding from the RPE monolayer. Furthermore, neighboring cells in these clusters exhibited immunoreactivity for several genes from different neuronal lineages. These structures are reminiscent of the recently reported ectopic “mini-retina” thought to originate from RPE after retinal damage in *Xenopus* tadpoles [43].

The genesis of the domes after ASCL1-NEUROD1 overexpression is not clear. We propose two hypotheses: (1) When monitoring the reprogramming process, we noticed that initially, many infected cells seem to be elongating and spreading out. That, together with the notion that the clusters are of varying size and do not always have the same cellular composition, may imply an aggregation process. In this model either cells would seek each other out and clump together or the proximity of cells itself might benefit survival and differentiation. (2) More intriguingly, as the infection of our culture is relatively sparse, it is possible that each cluster arises from one individual founder cell that is converted into a progenitor-like state. It then goes through a few rounds of cell divisions before the daughter cells exit the cell cycle and differentiate. Supporting this theory is the finding that when we started EdU treatment, only a few days after infection, most of the reprogrammed cells were already post-mitotic while surrounded by proliferative RPE.

### Limitations of this study

A caveat of the present study is that we used fetal RPE, which is highly proliferative and arguably closer to an immature/progenitor state. It might be that only a subpopulation of cells is preferentially infected or is permissive for reprogramming. It has been suggested that even adult RPE shows some degree of heterogeneity [16] and in the present work we did not strictly differentiate between retinal regions. Additionally, previous studies showed that the translation between model systems (2D vs organoid/*in vivo*; fetal vs. adult) and applied vectors can lead to varying outcomes on reprogramming [11], [17], [35], [36], [38], [44]. Further studies are necessary to assess the therapeutic potential of RPE with some focus on cell-type specific targeting and intraocular vector delivery. However, studies with iPSC-derived GMP-grade RPE for cell replacement are well underway [45], [46] and can be used as a vehicle to deliver reprogrammable cells to a damaged retina.

## Materials and Methods

### Primary cell culture

Human fetal eyes of gestational ages between 70 to 150 days were received from the Birth Defects Research Laboratory at the University of Washington following the approved protocol (UW5R24HD000836). After the removal of cornea, lens and retina, the remaining tissue consisting of sclera, choroid and RPE was incubated in 0.1 U/µl Collagenase IV (Gibco, 17104019) in HBSS (Gibco, 14025134) at 37 °C for 10 min. RPE was carefully peeled away in coherent sheets from the choroid making sure to avoid carryover of vasculature. RPE sheets were incubated for 15 min in 0.2 U/µl Collagenase IV in HBSS at 37 °C for 10-15 min and subsequently titruated with a P1000 pipette until a suspension of small clumps of ∼ 20 cells was achieved. Cells were spun down for 5 min. at 300g, supernatant was removed and pellet resuspended in prewarmed retinal differentiation medium (RDM). Cells were seeded into wells of a 24 well plate (1 ml/well) containing Matrigel (Corning, 354234) coated coverslips. RDM consisted in general of 50% DMEM (Gibco, 12430062), 50% DMEM/F12 (Gibco, 11330032), 1% antibiotic/antimycotic (Millipore Sigma, A5955-100ML), 1% NEAA (Gibco, 11140050), 1-10% FBS (Corning, 35-011-CV), 2% B27 (Gibco, 17504044).

For the first experiment (Fig. 2) cells were isolated from a donor of gestational day 137 (D137) and cultured in RDM with 10% FBS until processing for scRNAseq or staining. For all following experiments we used RDM with 5 % dialyzed FBS (Gibco, 26400044) and B27 without Vitamin A (Gibco, 12587010) from plating until the day of infection when we reduced the FBS concentration to 1%.

The screen (Fig. 3) was performed on RPE derived from D91 for control and library samples, which were cultured for 9 days before infection and for 11 further days before scRNAseq. At the same time an additional sample from D147 was infected at DIV8 and harvested at DPI18.

CMV>ASCL1-NEUROD1 overexpression was performed on a D115 sample, which was infected at DIV8 and harvested for scRNAseq (Fig. 4) or immunostaining (Fig. 5) on DIV28. After infection, half the wells were treated with 50 ng/ml bFGF (R&D Systems, 3718-FB-025) and 10 µM Activin A inhibitor (SB431542 Gibco, 120-14E-10UG).

For cell harvesting, media was removed and wells were washed 1x with DPBS and then incubated for 10 min at 37°C with TrypLE (Gibco, 12605028). The reaction was stopped by adding 1 ml of medium and the suspension was transferred to a fresh tube. Cell clumps were disrupted by pipette mixing with a P1000 pipette and passing through a 70 µm strainer.

### Immunostaining

Cells cultured on glass coverslips were washed 1x with DPBS and fixed with 4 % PFA for 15 min at room temperature followed by 3 more washes with PBS. For permeabilization and blocking, coverslips were incubated in 0.5 % Triton X-100/2 % normal horse serum (Vector Laboratories, S-2000-20) for 30 min at room temperature. After 3 washes with PBS, primary antibody in PBS was applied to the coverslips and incubated for 1.5 h at room temperature. Subsequently samples were washed 3 times in PBS and incubated in secondary antibody and DAPI for 1.5 h before mounting on a microscope slide.

EdU detection was performed using EdU ClickIT kit (Invitrogen, C10340) according to manufacturer’s protocol after permeabilization and before the primary antibody.

**Table 1:** Primary antibodies used in this study.

| Antibody | Dilution | host species | part number | company |
|---|---|---|---|---|
| ASCL1/MASH1 | 1:200 | mouse | BDB556604 | BD Bioscience |
| ASCL1/MASH1 | 1:200 | rabbit | ab211327 | Abcam |
| DCX | 1:500 | goat | sc-8066 | Santa Cruz |
| GFP | 1:300 | chicken | ab13970 | Abcam |
| HuC/D | 1:300 | mouse | A-21271 | Invitrogen |
| MKI67 | 1:200 | rabbit | BDB550609 | BD Bioscience |
| OTX2 | 1:300 | goat | AF1979 | R&D Systems |
| PMEL | 1:500 | mouse | 38815S | Cell Signaling |
| RCVRN | 1:200 | rabbit | AB5585-I | Millipore Sigma |
| TUBB3 | 1:500 | mouse | 801202 | BioLegend |
| TUBB3 | 1:500 | rabbit | 802001 | BioLegend |
| VSX1 | 1:200 | rabbit | 23566-1-AP | Proteintech |
| VSX2 | 1:200 | mouse | sc-365519 | Santa Cruz |
| ZO1 | 1:500 | rabbit | 61-7300 | Invitrogen |

### Determination of proliferation kinetics

Cells from three different donors were cultured on coverslips for up to 21 days. Coverslips for each donor were fixed at several intervals between 2 and 19 days and stained for MKI67. For image analysis we first used a custom Fiji [47] macro to convert czi format pictures to tif and to split channels before segmentation and quantification with CellProfiler [48]. In brief, we first segmented all nuclei via the DAPI signal and then identified proliferating cells in the MKI67 channel. We then related the resulting objects to yield MKI67-positive nuclei and calculated their percentage relative to all nuclei. The total cell density was calculated as the number of nuclei per scanfield from the same images.

### FACS sorting

After harvesting the D115 sample, FACS enrichment for mScarlet positive single cells was performed at the Department of Laboratory Medicine and Pathology Flow Cytometry Core (CC101467) at the University of Washington. RPE cells were harvested as previously described and passed through a cell strainer to obtain a single-cell suspension. Cell suspensions were sorted using the SymphonyS6 cell sorter (BD Biosciences). Live cells were gated based on their Forward and Side Scatter profiles and singlets were gated based on a bivariate plot of Forward Scatter Height versus Forward Scatter Area. The direct mScarlet signal in RPE cells across conditions was detected using the BYG584-P/PE laser. Uninfected cells were used as a gating control and 3-way purity settings were used to yield between 50000-100000 cells from each sample. Collection tubes were coated with FBS and filled with 50 µl PBS.

### Constructs

For the multiplexed screen we first designed a shared lentiviral backbone containing the expression cassette of mScarlet3 [49] driven by the CMV promoter. Adjacent to the CMV, we placed an EF1a promoter facing in the opposite direction on the antisense strand followed by the insertion site and a polyadenylation signal. This backbone was used to insert either one transgene or two transgenes separated by selfcleaving peptides (P2AT2A) followed by a constant sequence and a unique, construct specific barcode. To allow successful demultiplexing and account for possible sequencing errors, we used the R package DNAbarcodes [50] to design 10 bp long barcodes with a maximal Levenshtein distance of 3. At Vectorbuilder, plasmids were cloned separately but pooled in equimolar ratio before generating lentiviral particles to yield the final combined library. All constructs were acquired from VectorBuilder and are summarized in table 2.

**Table 2:** Summary of lentiviral vectors.

| Vectorbuilder ID | overexpression cassette | barcode |
|---|---|---|
| VB230117-1321vkj | CMV>ASCL1-TuGFP | - |
| VB230112-1267zah | pHES1>eGFP | - |
| VB230112-1267zah | pHES1>ASCL1-eGFP | - |
| VB250206-1630veh | Ef1a>ASCL1;CMV>NLS.mScarlet3 | GGAAGAAGAA |
| VB250206-1631mkr | Ef1a>ASCL1-EZH2;CMV>NLS.mScarlet3 | CAGAGAAGAA |
| VB250206-1632awa | Ef1a>ASCL1-LMO1;CMV>NLS.mScarlet3 | ACCAGAAGAA |
| VB250206-1634hdg | Ef1a>ASCL1-NEUROD1;CMV>NLS.mScarlet3 | GACGGAAGAA |
| VB250206-1635qve | Ef1a>ASCL-RAX2;CMV>NLS.mScarlet3 | AGTGGAAGAA |
| VB250206-1636ydf | Ef1a>NEUROD1-ETV1;CMV>NLS.mScarlet3 | TCACGAAGAA |
| VB250206-1638zyg | Ef1a>RAX2;CMV>NLS.mScarlet3 | CGCCGAAGAA |
| VB240318-1074fmh | CMV>NLS.mScarlet3 (ctrl) | CCGTCTCACT |
| VB250930-1548dgm | CMV>ASCL1-NEUROD1-mScarlet3 | - |
| VB250930-1550sjn | CMV>mScarlet3 (ctrl) | - |

### Data mining

ATAC seq raw data of adult and fetal RPE ([26], GSE126847) was kindly provided by Dr. Kelly Frazer. Read preprocessing, alignment and peak calling was performed using the ATAC pipeline from ENCODE [51].

From single cell ATAC data on human fetal retina [23], we used the fragments file produced by cellranger atac to filter fragments originating from photoreceptors and bipolar cells into individual files to create a pseudobulk representation. Macs2 [52], [53] was used to call peaks on these. Bedops [54] was used for peak operations and HOMER [55] for peak annotation and motif binding prediction.

### Single cell analysis

Single cell RNAseq libraries were prepared using 10x Genomics’ Chromium Next GEM Single Cell 3’ Reagent Kit v3.1 (10x Genomics, PN-1000268) according to manufacturer’s protocol. Samples were submitted for short read sequencing either at Seattle Genomics service core (NextSeq) or Northwest Genomics Center (NovaSeq) both at the University of Washington. Cellranger versions 7.2.0 or 9.0.0 were used for the processing of sequenced data. If necessary, fastq files were generated from raw sequencer output using cellranger mkfastq. A custom genome reference was created via cellranger mkref by adding the sequences of fluorescent reporters as well as parts of the lentiviral genome to human GRCh38 (2020-A GENCODE v32/Ensembl98; https://www.10xgenomics.com/support/software/cell-ranger/downloads).

Cellranger count was used for read alignment and counting. All analyses of the resulting count matrices and plotting of results was done in R [56] using the Seurat package [57].

### Screen demultiplexing

As the used indexing strategy with the Dual Index Kit TT, Set A (10x Genomics, PN-1000215/PN-3000431) employs unique indices, each sample can be identified by the presence of either i5 or i7 index. To maximize the number of sequencing reads for the analysis we therefore first looked at reads that were not assigned to any sample (Undetermined) and searched for the presence of either i5 or i7 index and assigned them to the respective sample using fastq-multx [58]. We combined these reads with the original, sample specific fastq files for the downstream analysis.

For demultiplexing we used a pipeline inspired by CellTag [59]. It is based on the fact that bam files that are output by cellranger contain aligned reads tagged with the cell barcode. Additionally, as the construct barcodes are transcribed from the same locus as the transgene, they are present on the same mRNA molecule and are captured and sequenced together. Using samtools [60] we looked for 10 bp of the constant pattern to detect reads in the bam file that contain a construct barcode. Next, we used the celltag.parse.reads.10x.sh script from CellTag to parse the reads yielding a table with a mapping of read sequence to cell barcode. Final cell to construct barcode mapping was achieved with an R script using the demultiplex function of the DNAbarcodes package with metric set to “seqlev” (Levenshtein) and distance <3. This output was then added as meta data when creating the corresponding Seurat objects.

## Data availability

All sequencing data produced in this publication was uploaded to Gene Expression Omnibus with the accession number GSE328663.

## Acknowledgements

We would like to thank Dr. Davide Ortolan for providing access to the Reshape 2 software, and Dr. Kelly Frazer for the ATAC seq raw data of adult and fetal RPE. Catherine Ray provided excellent technical assistance. We would like to also thank the members of the BDRL (Birth Defect Research Laboratory): Ian Glass, Kimberly A. Aldinger, Dan Doherty, Ian G. Phelps, Jennifer C. Dempsey, Kevin J. Lee, and Lucinda A, for their help with the human retinal tissues.

## Author contributions

LK: conceptualization, experimental design and execution, bioinformatic analysis, interpretation of data, writing of manuscript

ALG: tissue dissection, tissue culture, staining

JP: tissue culture, staining

MP: FACS, editing of the manuscript

JW: tissue dissection, scRNAseq, editing of the manuscript

TAR: conceptualization, funding acquisition, editing of the manuscript

## Supplementary Figure Legends

Supplementary Figure 1: A) Heatmap of RPE genes, markers for fibroblasts and epithelial to mesenchymal transition. Reprogrammed cells retain residual RPE expression pattern and are closer related to RPE than to fibroblasts. B) Differentially expressed genes between RPE, NPre and neur clusters.

Supplementary Figure 2: A) From human fetal single cell transcriptomic data we found pseudotime trajectories leading from retinal progenitor cells to photoreceptors. We identified genes whose expression change along that axis. RPC: retinal progenitor cells, AC: Amacrine cells, BP: Bipolar cells, PR: Photoreceptors. B) Genes from A) were additionally evaluated by looking whether they are upregulated during FGF-mediated RPE transdifferentiation in chicken. C) From single cell ATAC sequencing of human fetal retina and bulk ATAC sequencing of RPE we found peaks of open chromatin that contained OTX2 binding motifs. We then identified peaks that are common to RPE and photoreceptors (PR) but not bipolar cells (BP). These peaks were further analyzed for enriched binding motifs of other transcription factors. D) ASCL1 expression is highest in the clusters originating from cells that received the library with reprogramming vectors Npre, neur. NEUROD1 expression causes a shift in expression thus creating a smaller, distinct cluster of reprogrammed cells. MKI67 marks proliferating cells and NES is highly expressed in clusters of reprogrammed cells. E). Heatmap with RPE, fibroblast and EMT markers indicating the RPE origin of the reprogrammed cells.

Supplementary Figure 3: A) Overexpression of ASCL1 and NEUROD1 in human fetal RPE (D103) results in mScarlet positive clusters 10 days post infection even without FA treatment. These cells are also positive for early neuronal markers DCX and/or MAP2 albeit without prominent arborization. B) UMAP of integrated D59/76 fetal human retina used for reference mapping.

Supplementary Figure 4: A) Overexpression of ASCL1 and NEUROD1 and treatment with FGF and AAI in human fetal RPE results in mScarlet positive clusters with vast DCX-positive arborization. B) Top: Transduction of cultured RPE with the CMV>ASCL1-NEUROD1 lentivirus produces neurogenic clusters even in the absence of FA treatment. Bottom: Transduction with the control virus does not produce DCX positive cells nor mScarlet positive clusters even with the addition of FA. C/D) Control virus infection and FA treatment does not produce cells positive for cell type specific markers VSX1, HuC/D (C) or RCVRN, VSX2 (D). Scale bars 50 µm.

