## Supplementary figures and images for "Stimulating neurogenesis of distinct retinal lineages in human retinal pigmented epithelium (RPE) with proneural transcription factors"

### Supplemental Figure 1

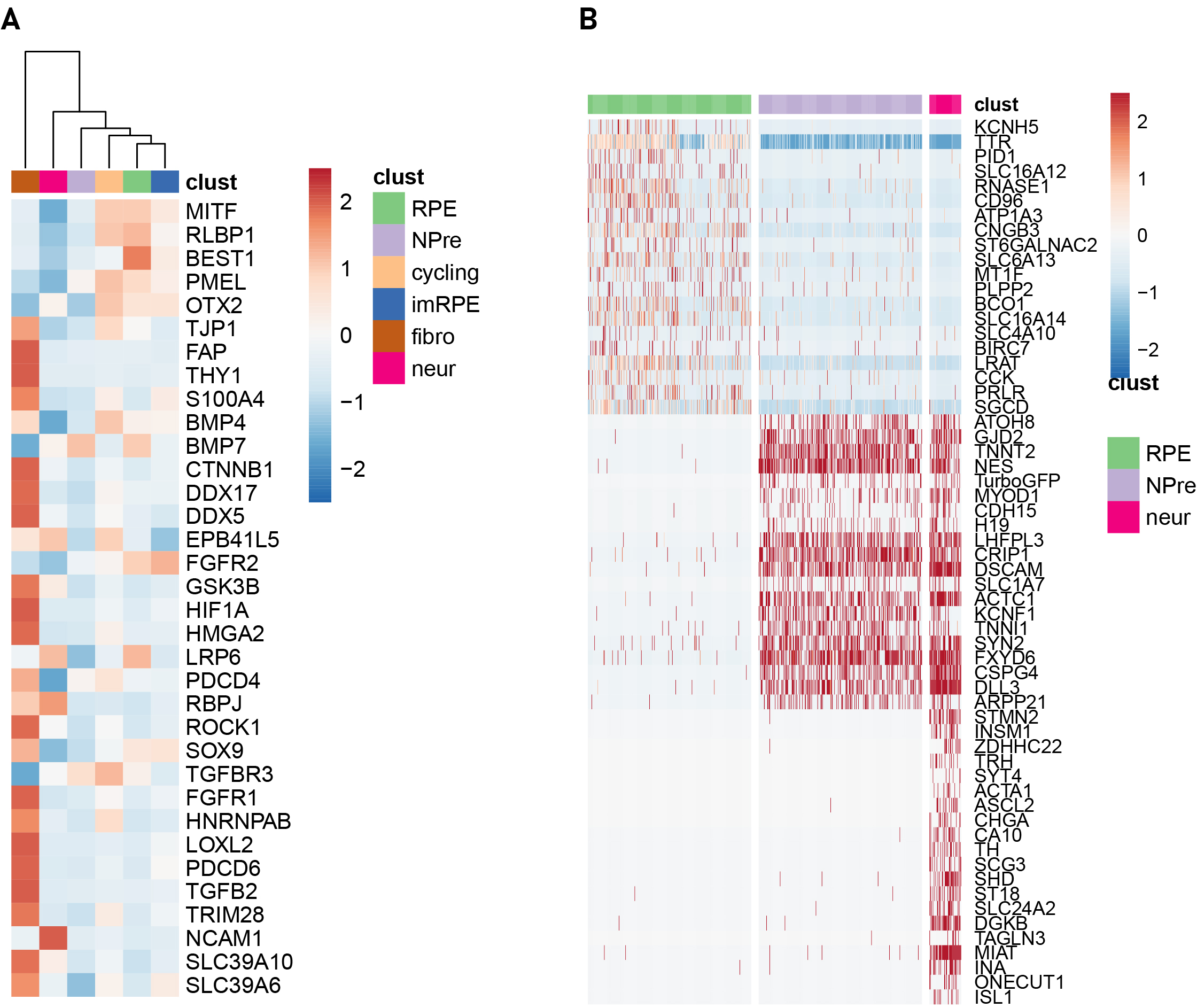

### Supplemental Figure 2

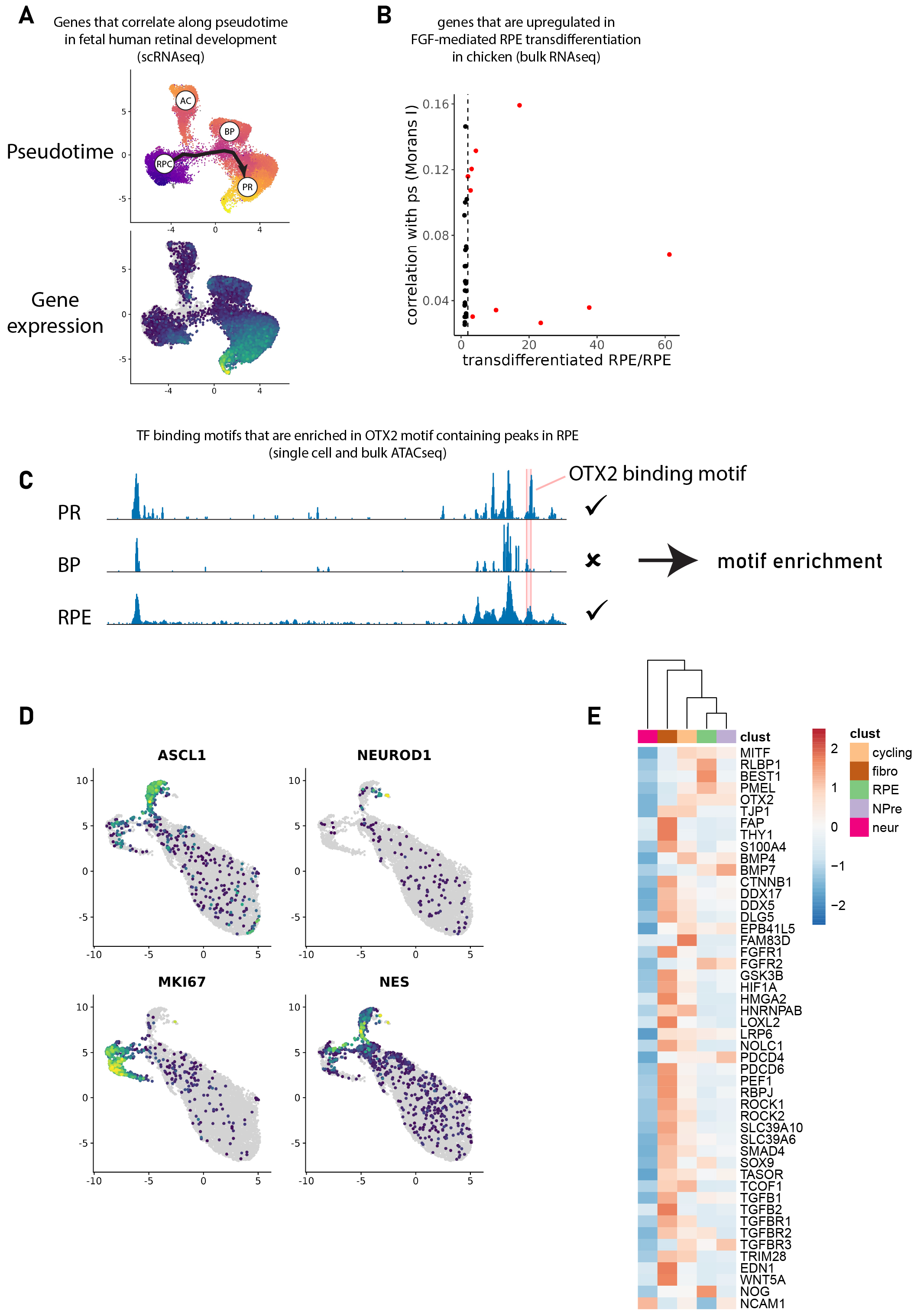

### Supplemental Figure 3

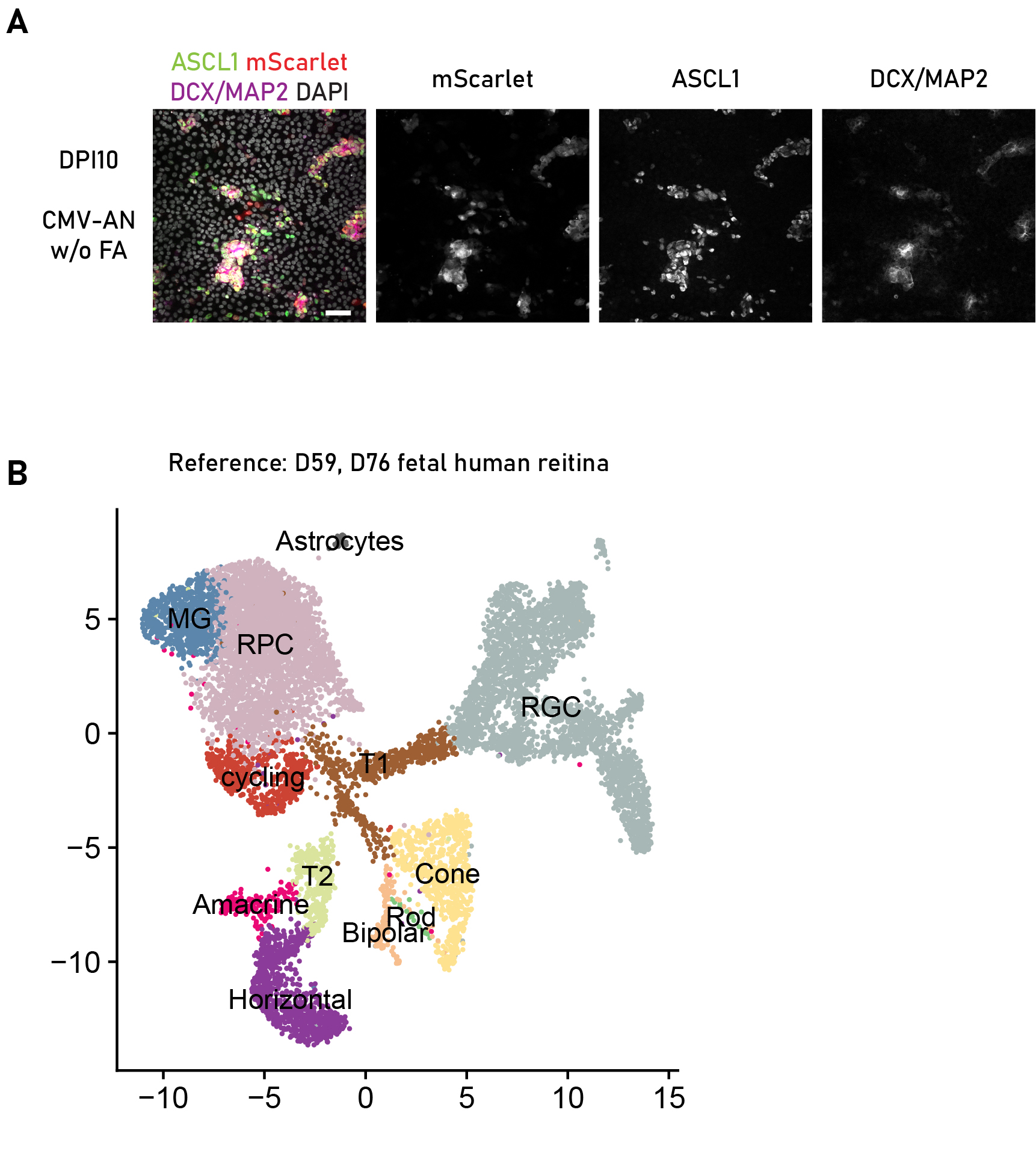

### Supplemental Figure 4

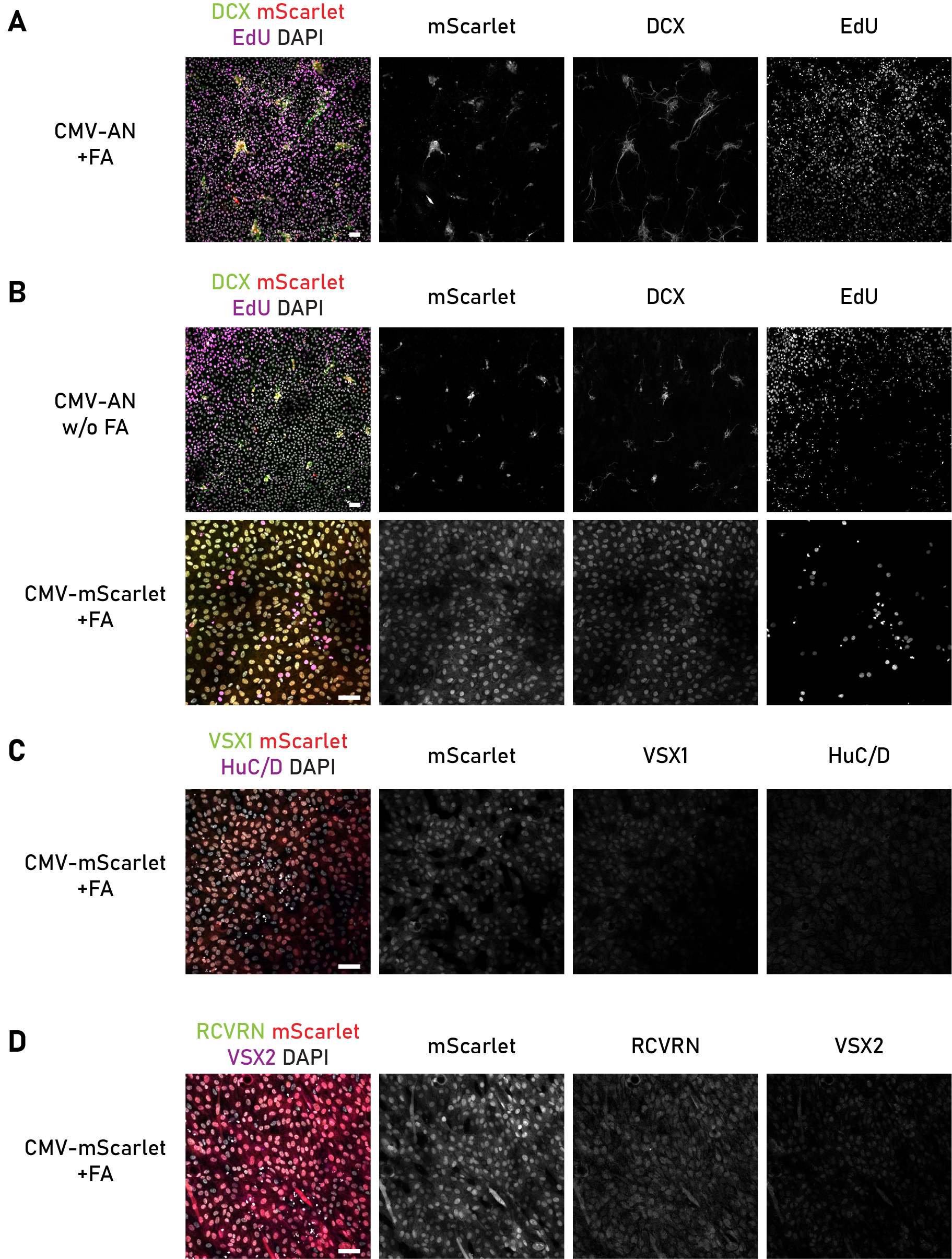
